# Left-DLPFC theta-burst stimulation produces delayed effects on musical pleasure without affecting piano-melody learning

**DOI:** 10.64898/2026.09.20.753042

**Authors:** Sofia Simonetto-Rizk, Yanan Liu, Jérémie Ginzburg, Robert J. Zatorre, Emily B. J. Coffey

**Author notes:** **Corresponding author:** [Sofia Simonetto-Rizk, ].

## Abstract

Reward operates as a cycle in which a stimulus produces a pleasure response, and the degree of pleasure experienced reinforces the behaviours that lead back to it. Music provides a useful model for examining this cycle because it is intrinsically pleasurable, can be learned, and engages reward-related frontostriatal circuitry that overlaps with circuitry involved in primary and secondary rewards, such as food, sex, and money. Transcranial magnetic stimulation (TMS), a non-invasive stimulation method, has been shown to produce bidirectional effects on music-evoked pleasure by increasing or decreasing activity in cortical regions, such as the left dorsolateral prefrontal cortex (DLPFC), that are functionally connected to subcortical regions, including the striatum. Here, we further characterize the reward cycle by incorporating TMS into a paradigm that combines musical pleasure ratings with an early musical skill acquisition task. In a within-subjects, counterbalanced crossover design, 23 participants with 2-10 years of musical experience received intermittent TBS (iTBS; excitatory) and continuous TBS (cTBS; inhibitory) over the left DLPFC in separate sessions. Participants rated the pleasantness of short piano melodies before stimulation (T0), immediately afterwards (T1), and 24 hours later (T2). After stimulation, they learned to play the melodies, and pitch and rhythm accuracy were assessed during three performance phases, namely acquisition, immediately after acquisition, and 24 hours later. Liking did not differ between conditions at T1; however, the increase in liking from baseline (T0) to T2 was significantly larger following iTBS than cTBS, a divergence that emerged across the 24-hour interval spanning both learning and sleep. Pitch and rhythm accuracy did not differ between conditions at any phase, despite improvement across trials in both. These results indicate that TMS to the left DLPFC can produce delayed, condition-dependent shifts in musical liking without a parallel change in early skill acquisition.

## I. Introduction

Reward is a natural process by which the brain associates stimuli (substances, situations, events, or activities) with a positive or desirable outcome (Lewis et al., 2022). The accompanying experience of pleasure often motivates behaviour. Reward is traditionally described as a cycle with two linked components: contact with a stimulus produces a consummatory hedonic response, or ‘liking’, and the experience of that response is encoded through learning, which increases the probability that the behaviours leading back to the stimulus will recur (Berridge & Kringelbach, 2008). Reward is therefore best understood as the mechanism that orients learning toward reward-producing behaviour.

Music is an interesting stimulus in this regard, as it is both intrinsically pleasurable and one that can be learned through production (Zatorre, 2015). Past research on reward focused on the hedonic response as a product of satisfying homeostatic needs such as hunger, sex, and bodily comfort, as well as avoiding pain (Reybrouck & Eerola, 2022). This response was associated with both primary rewards, like food and sex, and secondary rewards, like money, and the reward circuitry supporting it primarily involved fronto-striatal projections of dopamine, spanning both cortical and subcortical regions of the brain (Sescousse et al., 2013). However, music, an abstract aesthetic reward, possesses the unique ability to elicit pleasure similar to primary and secondary rewards through this same mesolimbic circuitry, without satisfying any homeostatic need (Blood & Zatorre, 2001).

Neuroimaging studies have since confirmed this striatal engagement (Salimpoor et al., 2013; Zatorre & Salimpoor, 2013), and two studies using PET have shown that the response involves dopamine directly: Salimpoor et al. (2011) found dopamine release in the caudate during anticipation of a peak-pleasure moment and in the nucleus accumbens during the peak hedonic response itself, and Fritz et al. (2026) more recently showed that pleasant music engages the phasic dopaminergic D1-receptor system. This evidence remains correlational, however, and establishing that dopamine does more than accompany musical pleasure requires intervening on the system directly. Ferreri et al. (2019) did so pharmacologically, manipulating synaptic dopamine availability with levodopa and showing that this shifted the pleasure participants reported while listening to music, which established a causal role for dopaminergic signalling in musical pleasure. A follow-up study in the same cohort showed that this causal link extends into memory: excerpts rated as more pleasurable under placebo were better remembered a day later, and disrupting dopaminergic transmission with risperidone weakened this relationship, specifically for recollection rather than for familiarity-based recognition (Ferreri et al., 2021).

This causal evidence, however, remains limited in scope. Pharmacological interventions involve ingesting a substance that acts systemically, so its effects are diffuse rather than anatomically specific, and the causal claim they support concerns dopaminergic transmission broadly rather than the reward circuitry underlying musical pleasure in particular. In this sense, the precision of the claim is limited in much the same way that neuroimaging limits it, only from the opposite direction: imaging localizes the response without establishing that it is causal, and pharmacology demonstrates that it is required without allowing for localization. Transcranial Magnetic Stimulation (TMS), a non-invasive brain stimulation technique, can be used to target the reward circuitry underlying musical pleasure directly via the direct stimulation of a cortical region functionally connected to the striatum. Mas-Herrero et al., (2018) used intermittent theta-burst stimulation (TBS) over the left DLPFC, at coordinates previously shown to induce striatal dopamine release (Strafella et al., 2001; see also Keck et al., 2002; Kanno et al., 2004), in seventeen participants without musical training. Across three sessions at least 24 hours apart, non-musician participants received intermittent TBS (iTBS), which increases cortical excitability, continuous TBS (cTBS), which decreases it (Huang et al., 2005), or sham stimulation. They then listened to fifteen naturalistic 45-second excerpts, five self-selected and ten experimenter-selected, while rating their pleasure in real time and while physiology was recorded. iTBS increased both measures and cTBS decreased them, regardless of whether music was self-selected or experimenter selected. A follow-up fMRI study replicated and localized these effects: excitatory stimulation enhanced and inhibitory stimulation disrupted striatal responsiveness to hedonic value, and changes in the ventral striatum, particularly the nucleus accumbens, predicted changes in reported pleasure, with no change at the stimulation site itself (Mas-Herrero et al., 2021).

A further limitation concerns what is actually measured: in Ferreri et al., (2019, 2021), the outcome is memory for an excerpt that was only listened to. The action step of the reward cycle, as operationalized here, describes goal-oriented behaviour that goes beyond recognizing a stimulus; it includes the motivation to practice and reproduce it. This step is captured by the learning process which spans acquisition, retention, and delayed retention (i.e., practice, recall and long-term retention). Learning a melody requires encoding its pitch and rhythmic structure, mapping that structure onto movements, and refining the mapping across attempts until the melody can be produced (Herholz & Zatorre, 2012). Listening and recognizing engage only the first of these stages. Recognition is supported by declarative memory, whereas production and practice are thought to depend on procedural systems (Squire, 2004); the two systems rely on different brain structures and are consolidated through partly shared sleep mechanisms (Boutin & Doyon, 2020). A finding that reward shapes one (Ferreri et al., 2021) therefore does not entail that it shapes the other. Thus, it remains an open question whether a shift in hedonic response changes only how music is experienced or also how quickly it is acquired and whether it is retained and retrieved over time. Answering this question requires a way to act directly on the circuitry underlying musical pleasure, which TMS provides, and to follow its consequences beyond what is remembered, into what can be executed, over a time course long enough to capture lasting effects.

Together, these gaps raise three open questions about the relationship between musical pleasure, learning, and reward. First, does a difference in liking between excitatory and inhibitory stimulation appear while cortical excitability is altered, after it has returned to baseline, or both? Second, does learning follow the same pattern, with any effect of stimulation on acquisition present immediately or emerging across the following day? Third, does stimulation of reward circuitry produce measurable effects at all with melodies short and simple enough to be learned in a single session? We hypothesize (i) that iTBS would increase subjective liking ratings for short piano melodies and cTBS would decrease it; and (ii) that learning would follow the same pattern as liking: under the excitatory condition, melodies for which liking was enhanced would be performed better, and under the inhibitory condition, melodies would be performed worse.

To address these questions, the present study combines manipulation, time course, and rewarding stimuli in a within-subjects design. In counterbalanced sessions, participants received iTBS and cTBS over the left dorsolateral prefrontal cortex, rated short piano melodies for pleasantness, and then learned to play them, with both liking and learning assessed immediately after stimulation, while cortical excitability was still altered, and again 24 hours later, once it had returned to baseline. The 24-hour assessment matches the interval over which pharmacological manipulation of dopamine was previously shown to alter recollection (Ferreri et al., 2021); no study manipulating reward circuitry, pharmacologically or with TMS, has yet extended that interval to a measure of procedural skill rather than declarative memory. Melodies were designed to be short and of comparable difficulty to permit intermediate-level musicians to learn and retain a set of 20 melodies within a single session. In doing so, this design offers a first exploration of whether the perception and action steps of the reward cycle, the hedonic response to music and the learned behaviour that follows it, are shaped together by a common manipulation of reward circuitry, or whether they can be dissociated.

## I. Materials and Methods

### PARTICIPANTS

An a priori power analysis was conducted using G*Power to estimate the sample size required for a within-subjects repeated-measures analysis. The analysis was based on the effect size reported by Mas-Herrero et al. (effect size *f*=.33068), with statistical power (1-β error prob.) set to .8 according to standardized procedures (Cohen, 1992). The output indicated a minimum of 17 participants for a within-factor repeated-measures analysis. A final sample of 23 healthy participants (10 male, M =23.03 ± 4.52 years) was included in the behavioral analyses accounting for participant dropout and technical malfunctions. All participants provided informed consent before taking part in the study. The experimental protocol was approved by Concordia University’s Human Research Ethics Committee and the Montreal Neurological Institute Ethics Review Board.

Participant screening and characterization were conducted using three questionnaires: the Montreal Music History Questionnaire (MMHQ; Coffey et al., 2011), to screen for number of years and type of musical experience, a TMS Adult Safety Screening Questionnaire, the Barcelona Music Reward Questionnaire (BMRQ), to screen for threshold level of musical reward sensitivity (≥65) (Mas-Herrero et al., 2013). In order to be eligible for the study, participants had to be right-handed, have normal hearing, have between 2 and 10 years of musical experience, and have no contraindications for the safe implementation of TMS protocols. Participants with previous piano experience were excluded because the study required participants to learn unfamiliar piano melodies to characterize early musical training. Participants taking medications that target the central nervous system, such as antidepressants or sleeping medication, those who had a documented history of significant neurological, psychiatric, or sleep disorders, and have consumed recreational drugs or alcohol within 24 hours of their scheduled participation time were excluded.

### EXPERIMENTAL DESIGN

#### Procedure

The study used a within-subjects, randomized, counterbalanced crossover design. Each participant completed two experimental blocks, one involving intermittent theta-burst stimulation (iTBS) and the other involving continuous theta-burst stimulation (cTBS) over the left dorsolateral prefrontal cortex. The order of stimulation conditions and the assignment of the two melody sets to stimulation conditions were counterbalanced across participants. Participants first completed the pre-stimulation liking-rating task (T0), which lasted approximately 5 to 10 minutes. Following T0 assessment, participants received one of the two stimulation paradigms (cTBS or iTBS).

Immediately after stimulation, participants completed the liking-rating task again (T1), followed by the piano-learning task. The learning task consisted of six acquisition trials and two immediate retention trials for each of the 20 melodies assigned to that experimental block. These tasks lasted approximately 45 minutes in total following the TMS stimulation. Participants returned 24 hours following each stimulation session for the delayed assessment. This session consisted of both the liking-rating task (T2), a physiological recording, and two delayed retention trials for each of the 20 melodies learned during the preceding block. This delayed session lasted a total of 45 minutes. The second experimental block followed the same procedure using the opposite stimulation condition and the other 20-melody set. The second block began at least 1 week after the completion of the delayed assessment from the first block to reduce potential carry-over effects.

### STIMULI

#### Musical stimuli

Forty piano melodies (2-5 s in duration, C major scale, 4/4) were selected from a larger pool of 70 candidate melodies following pilot testing. The 40 highest-rated melodies were divided equally into two 20-melody sets, namely V1 and V2. Each participant completed one experimental block with V1 and the other block with V2; thus, each participant rated and learned all 40 melodies across the two experimental blocks. The assignment of melody set to stimulation condition and the order of stimulation conditions were counterbalanced across participants. The order of melody presentation within each block was randomized. Pilot testing confirmed that the two sets did not differ significantly in pleasantness ratings or in playability for intermediate musicians within the constraints of the protocol.

### APPARATUS

#### TMS setup

A 70 mm figure-of-eight coil, Super Rapid Biphasic Stimulator, was used to administer the Transcranial Magnetic Stimulation (Magstim, Whitland, UK). A real-time optically tracked frameless stereotaxic system (Brainsight Frameless, Rogue Research Inc.) using an infrared camera for online coil positioning (Polaris Spectra, NDI) was used for subject tracking and coil positioning.

#### Active motor thresholding

During the first stimulation session, the active motor threshold (aMT) was estimated to establish the individualized signal strength for the following stimulation sessions. To determine aMT, the left primary motor cortex was stimulated using a single-pulse protocol to elicit motor-evoked potentials (MEPs) from the first dorsal interosseous (FDI; index finger). To identify the MT hotspot, we began with 50% of the highest maximum stimulator output, which yielded an MEP response for most participants. The optimal spot, i.e., where an MEP response in the FDI was observed, was found by moving the TMS coil in 0.5-1 cm increments on the scalp, starting from approximately 5cm lateral to the vertex of the head, applying one pulse per spot.

Intensity was increased in increments of 5% until a response was elicited. Once found, the aMT was defined as the minimum intensity that produced a visible MEP (> 200 μV) in 50% of 10 trials during isometric contraction of the target muscle. This threshold was then used to set the intensity of the stimulation (80% of aMT; in replication of Mas-Herrero et al., 2020)

#### Beam method

The left DLPFC stimulation site was determined during the first session using the BeamF3 method (Mir-Moghtadaei et al., 2015), an individualized scalp-based algorithm requiring three cranial measurements, based on the international 10-20 EEG system. The three measurements include the nasion-inion distance (NI), the left tragus-right tragus distance through the scalp vertex (TrTr), and the head circumference (HC) measured through the FPz-Oz plane of the international 10-20 EEG system. These measurements were used to calculate two coordinates, X and Y, which correspond to the arc length from the cranial midline and the radial distance from the vertex, respectively. The intersection of the two coordinates represented the scalp location overlying the left DLPFC (Mir-Moghtadaei et al., 2015). Once identified, the stimulation target was marked directly on the TMS cap. A real-time optically tracked frameless stereotaxic system (Brainsight Frameless, Rogue Research Inc.) was then used to align the BeamF3 target with the Brainsight platform and guide the coil positioning during stimulation sessions. An infrared camera (Polaris, NDI) provided continuous online monitoring of participant head position and coil placement throughout stimulation, ensuring accurate and consistent targeting.

#### TBS

Theta-burst stimulation was applied using the Magstim Super Rapid stimulator during the two sessions. It was delivered using a TBS pattern consisting of 3 pulses at 50Hz repeated at 5Hz (Huang et al., 2005). For iTBS, cycles of 2 second trains of TBS (10 bursts repeated at 5 Hz, each consisting of 3 pulses, therefore 30 pulses) followed by 8 seconds of no stimulation were repeated until 600 pulses were delivered (20 cycles for a total of 192 s). In the cTBS protocol, a 40-second train of uninterrupted TBS was given (200 bursts repeated at 5 Hz, consisting of 3 pulses each, therefore 600 pulses in total).

#### Piano-learning platform

Piano-learning performance was assessed using an open-source Music Paradigm platform adapted from a longitudinal design developed by Herholz and colleagues (2012); a related adaptation of this paradigm appropriate for a single learning session was also used by Jourde et al. (2026). Participants learned to reproduce 20 short melodies per experimental block on a MIDI keyboard, using their left hand to increase the difficulty of the task and allow more room for motor-skill improvement, given that all participants were right-handed. Each melody was first played to participants, accompanied by a visual representation of the required keys on a keyboard diagram displayed on the computer monitor, before participants attempted to reproduce it. The learning session consisted of six trials per melody. Visual cues, in which the key sequence was highlighted during melody presentation, were offered only on the second and fourth trials, to enable participants to learn the melodies; cues were withheld on the remaining trials to discourage reliance on them and to encourage auditory-motor, rather than visual-motor, learning. Only the four unguided trials (1, 3, 5, and 6) therefore contributed to the primary acquisition outcome (see Measures). After each of the six learning trials, participants received feedback on their pitch and rhythmic accuracy in the form of two coloured visual indicators: pitch feedback was binary (all keys correct or not), and rhythm feedback was given in three ranges of accuracy (good, moderate, poor). Immediately after the learning session, participants completed two retention trials without visual cues or feedback; the first (T7) was treated as a reorientation trial reflecting participants’ initial re-engagement with the unaided retrieval context, and the second (T8) served as the primary immediate-retention outcome. Approximately 24 hours later, participants completed two further retention trials under the same unaided conditions; as with the immediate-retention trials, the first (T9) was treated as a reorientation trial and the second (T10) served as the primary delayed-retention outcome.

### MEASURES

#### Subjective liking measure

Liking for each melody was assessed using a continuous rating scale implemented in PsychoPy. The scale ranged from (1) “No pleasure” to (7) “Very pleasurable”, with seven labeled tick marks providing reference points for the ratings. Participants rated the pleasantness of each melody at three time points: before stimulation (T0), immediately after stimulation (T1), and during the delayed assessment approximately 24 hours later (T2). Following the presentation of each melody, participants selected the rating that best represented their subjective level of pleasure.

#### Piano performance measure

The piano-learning task was selected as a controlled yet ecologically relevant model of complex auditory-motor learning, requiring the integration of auditory perception, pitch and temporal encoding, sequential motor planning, feedback, and memory (see Apparatus). Pitch and rhythm were scored separately to isolate distinct components of early musical learning across acquisition, immediate retention, and delayed retention. Pitch accuracy was calculated as the percentage of target notes reproduced correctly in the correct serial order, with the number of correctly reproduced notes divided by the total number of target notes in each melody.

Higher pitch-accuracy scores indicated better performance. Rhythm performance was quantified using a relative inter-onset interval (IOI) error metric. For each melody and trial, IOIs were calculated for both the reference melody and the participant’s performance. The absolute difference between corresponding IOIs was divided by the mean reference IOI, and the resulting relative errors were averaged across the melody and expressed as a percentage. Lower values indicated more accurate reproduction of the melody’s temporal structure. This approach allowed participants to perform a melody at a faster or slower overall tempo while receiving a high score when the relative rhythmic structure was preserved. Because rhythm-error scores were positively skewed, the raw percentage-error scores were transformed using a log(1 + x) transformation before statistical analysis.

### STATISTICAL ANALYSIS

#### General modeling approach

Statistical analyses were conducted primarily in Python using the “statsmodels” package (Seabold & Perktold, 2010), with the exception of the binomial generalized linear mixed-effects models used to analyze pitch accuracy, which were fit in R using the “lme4” package (Bates, Mächler, Bolker, & Walker, 2015). Linear mixed-effects models were estimated using restricted maximum likelihood, and the L-BFGS optimizer was used initially; models that failed to converge were refitted using the Powell optimizer. Pitch accuracy GLMMs were fitted using maximum likelihood (Laplace approximation) with lme4’s bobyqa optimizer. Stimulation condition was entered as a fixed effect, with iTBS (excitatory TMS) serving as the reference condition. The condition coefficient thus represented the difference between cTBS and iTBS. Model-specific random effects structures are described in the relevant analysis sections and account for repeated observations within participants, participant session blocks, and melodies, where applicable. Fixed effects were evaluated using Wald z-tests and two-sided 95% confidence intervals. Statistical significance was evaluated using α=.05. Separate Holm-corrected testing families were defined for liking, pitch performance, and rhythm performance. The liking family contained three tests. The pitch and rhythm families each contained three phase-specific stimulation-condition tests for acquisition, immediate retention, and delayed retention. These families were corrected separately rather than pooled across outcomes.

Effect sizes were reported alongside each model’s coefficients: Cohen’s *d* (the fixed-effect coefficient standardized by the model’s residual standard deviation) for continuous linear mixed-effects models, and odds ratios (the exponentiated coefficient) for binomial models. In a statistical summary table reporting the best-fit model’s condition effect at each learning phase, Cohen’s *U3* (Cohen, 1988) was additionally computed as Φ (d), interpreted as the proportion of the lower-scoring condition’s distribution falling below the higher-scoring condition’s mean. For pitch accuracy (binomial GLMM), the logit coefficient was first converted to a d-equivalent using the standard logit-to-*d*-equivalent using the standard logit-to-d transformation (d≈ *b* x 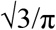; Chinn, 2000)

#### Assumptions

Model adequacy was assessed separately for the linear mixed-effects models and binomial generalized linear mixed-effects models. For all models, convergence status and estimated random-effect variance components were examined, and Model fit with random-effect variance components at or near zero was flagged as potentially singular (Bates, Mächler, Bolker, & Walker, 2015).

For linear mixed-effects models, the approximate normality and homoscedasticity of conditional residuals were assessed using conditional residual Q-Q plots, residual histograms, residual-versus-fitted plots, and residual distributions across stimulation conditions and a Spearman correlation between absolute residuals and fitted values. Shapiro-Wilk tests were used as supplementary screening measures for departures from normality and were not interpreted in isolation. Multicollinearity among fixed effects was assessed using variance inflation factors (VIF), with values greater than 5 flagged for further inspection.

For binomial GLMMs, normality and constant-variance assumptions for response-scale residuals were not imposed. Model adequacy was alternatively assessed using convergence status, random-effect variance estimates, Pearson-residual plots, and the Pearson dispersion statistic (Bolker et al., 2009). Dispersion statistics above 1.4 were flagged as potential overdispersion, whereas values below .70 were flagged as potential underdispersion. The conditional random-effect estimates were also examined for approximate normality using Shapiro-Wilk tests when the number of estimated random effects allowed for this assessment. Correlations among fixed-effect estimates were inspected using the model-based variance-covariance matrix, with pairs exceeding an absolute correlation of .70 flagged for further inspection, with particular caution applied to correlations involving interaction terms, which may in turn reflect the coding or scaling of the predictors rather than problematic model specification. For both model families, influential observations were identified using standardized conditional results where observations with an absolute standardized residual greater than 3 were examined for possible scoring or data-entry error and were retained unless an error was identified.

#### Liking analyses

The primary liking model predicted liking ratings from stimulation condition, timepoint (T0, T1, T2; T0 as reference), their interaction, and curriculum version, with the random-effects structure described above. A planned contrast tested whether the condition effect differed between T2 and T1. To control the family-wise error rate, the two condition x time interaction terms, the planned T2-versus-T1 contrast, and the condition x baseline-liking interaction were treated as a single three-test Holm-corrected family (Holm, 1979).

#### Learning analyses: outcome definitions

Pitch accuracy was coded at the target-note level as correct or incorrect, based on whether each note was reproduced at the correct pitch and serial position. It was summarized descriptively as the percentage of correctly reproduced target notes, with higher values indicating better performance. Rhythm performance was analyzed as relative inter-onset interval error, with lower values indicating more accurate temporal reproduction; because rhythm-error scores were positively skewed, they were transformed using log(1 + x) prior to model fitting. Curriculum was not included as a fixed effect in the learning models because curriculum assignment was counterbalanced with stimulation conditions across participants, decorrelating the two by design.

#### Learning analyses: Pitch

At each learning phase, condition effects were modeled three ways to assess model fit: a binomial GLM with cluster-robust standard errors, a binomial GLMM (lme4::glmer; Bates, Mächler, Bolker, & Walker, 2015), and a continuous LME on raw and logit-transformed percentage-correct, to identify the best-supported specification. AIC comparison and diagnostic checks favored the GLMM over the plain GLM at every phase; diagnostic checks favored the GLMM over both continuous LME specifications, which showed non-normal and heteroscedastic residuals at every phase (*see* Supplementary Materials). The GLMM was retained as the model of record for pitch accuracy. The acquisition-phase GLMM included centered trial number and its interaction with condition; immediate- and delayed-retention GLMMs modeled condition alone, with delayed retention additionally adjusting for T8 performance. Pitch mixed-model results were interpreted cautiously given evidence of ceiling effects in pitch accuracy.

#### Learning analyses: Rhythm

Acquisition-phase rhythm performance was modeled using a linear mixed-effects model including stimulation condition, centered linear trial number, centered quadratic trial number, and their interactions with stimulation conditions. The quadratic specification was retained to accommodate a decelerating learning curve because it provided a better fit than the corresponding linear specification according to AIC (*see* Supplementary Materials). Immediate- and delayed-retention rhythm performance phases used linear mixed models with condition as the sole fixed effect. The delayed-retention model is additionally adjusted for immediate-retention performance.

## II. Results

### Model diagnostics

Model diagnostics supported the reported specifications. For the five continuous linear mixed-effects models (liking and rhythm outcomes), conditional residuals were approximately normal and homoscedastic on visual inspection of Q-Q, histogram, and residual-versus-fitted plots; statistically significant Shapiro-Wilk results reflected the large sample sizes rather than substantive departures from normality, and the rhythm immediate-retention model satisfied the Shapiro-Wilk criterion outright (*p* = .18). For the three binomial pitch GLMMs, Pearson-residual plots showed the downward fan characteristic of near-ceiling accuracy, but model dispersion remained close to 1.0, indicating adequate fit; accordingly, pitch results were interpreted with caution regarding ceiling effects. Across all models, fewer than 4% of observations exceeded a standardized residual of ±3, and these were verified as valid extreme scores rather than data-entry errors and retained.

### Liking

Liking ratings did not differ significantly between conditions at baseline (*see* Table 3). Under Excitatory stimulation, mean liking ratings increased significantly from baseline (T0) to 24 hours Post-TMS (T2), and at the Condition x Time interaction level for the 24 hours Post-TMS timepoint (T2) (*see* Table 3), following multiple comparisons correction (Holm correction). This demonstrates that participants in the Inhibitory condition showed a significantly smaller liking increase than participants in the Excitatory condition (*see* Figure 1 & 2). Prior to and following multiple comparisons correction, no statistically significant change was found from the Pre-TMS baseline (T0) to the Post-TMS timepoint (T1), nor at the Condition x Time interaction level. This indicates that participants in either stimulation condition did not differ in how their liking changed from baseline to immediately after stimulation. However, a planned-contrast analysis between 24 hours Post-TMS (T2) and Post-TMS (T1) revealed that a divergence by condition emerged specifically over the 24-hour interval, such that the Excitatory group had a higher liking rating than the Inhibitory group. Curriculum version was non-significant, indicating that the two melody sets (v1 and v2) did not differ systematically in liking, which provided support for the validity of the counterbalancing curriculum assignment across conditions.

**Figure 1.**
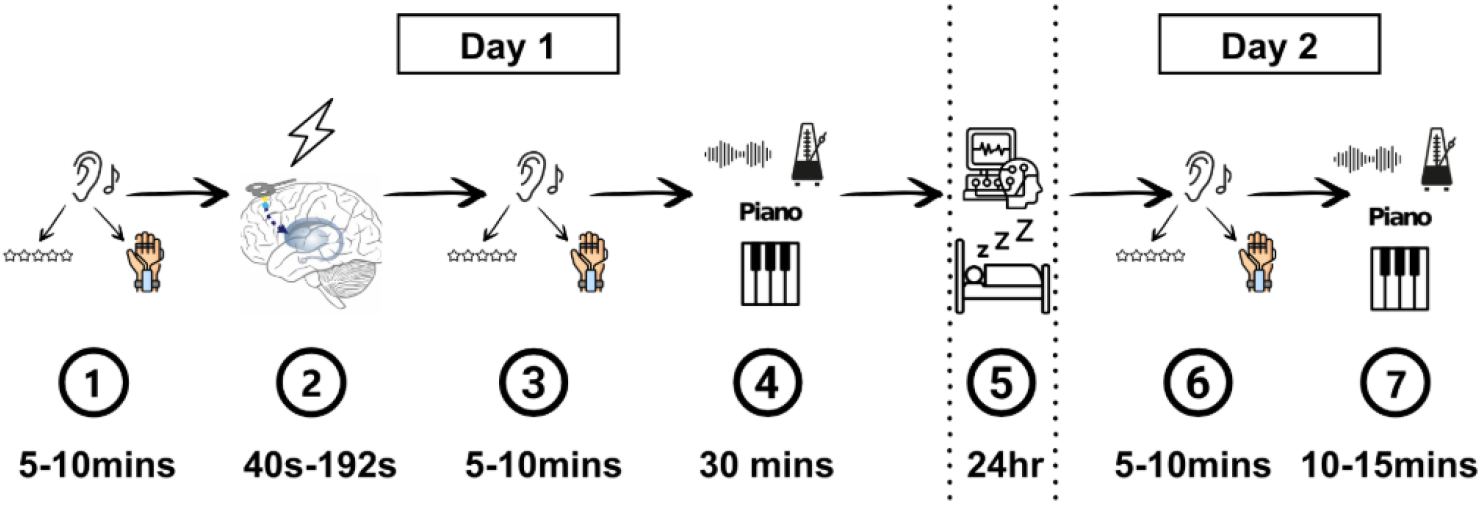
Timeline of a single experimental session across two days. Each participant completed this procedure twice, once with iTBS and once with cTBS in counterbalanced order, at least 24 hours apart. (1) Baseline listening and liking ratings of the 20 melodies (T0). (2) Theta-burst stimulation over the left DLPFC (cTBS: 40 s; iTBS: 192 s). (3) Post-stimulation listening and liking ratings (T1), completed while cortical excitability was still altered. (4) Piano-learning task involving six acquisition trials and two immediate-retention trials. (5) 24-hour interval, including sleep. (6) 24-hour post-stimulation listening and liking ratings (T2) once excitability has returned to baseline. (7) Piano-learning task involving two delayed retention trials. Durations are approximate.

**Figure 2.**
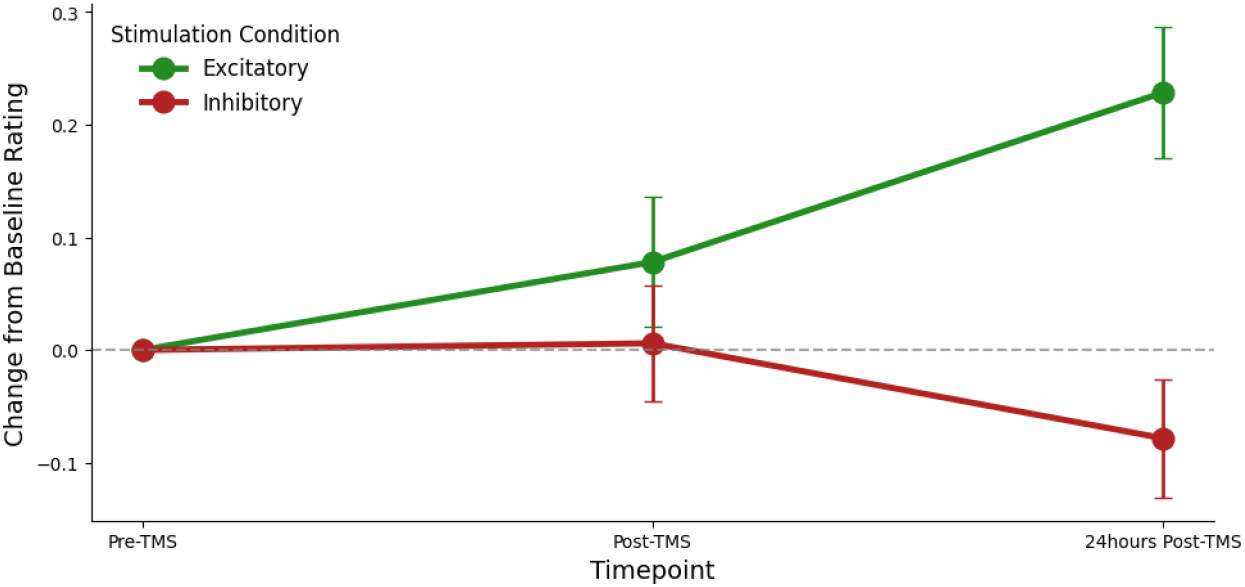
Change in liking relative to baseline by stimulation condition and timepoint. Points show the mean change in liking rating from baseline (T0) to immediately after stimulation (T1) and 24 hours later (T2) under excitatory (iTBS) and inhibitory (cTBS) stimulation. Liking was rated on a scale from 1 (no pleasure) to 7 (very pleasurable). Error bars show ±1 *SE*. The dashed horizontal line marks no change from baseline. *N*=23.

Effect sizes and Cohen’s U3 were examined for every term in the confirmatory family, standardized by one common residual SD applied consistently across all three contrasts, in alignment with the common standardizer (“*d*-with”) approach to standardized contrasts. The Condition x Time (T1) interaction showed near-complete overlap (*see* Table 4), consistent with its non-significant result and with no meaningful over- or under-representation of either condition extreme in the hour immediately following stimulation. The Condition by Time (T2) interaction showed more substantial separation (see Table 4), where, in this sample, twenty-four hours after stimulation, most Excitatory-condition melodies (roughly 65%) gained more in liking than the average Inhibitory-condition melody did. The planned T2-versus-T1 contrast showed a comparable degree of separation (*see* Table 4), consistent with this divergence emerging specifically over the 24-hour interval rather than being present immediately after stimulation. This means that the gap favoring Excitatory over Inhibitory in liking change was not just present at 24 hours, but reliably larger at that point than it had been right after stimulation, indicating the difference between conditions emerged sometime between the immediate and 24-hour assessments rather than being there at the beginning.

**Table 4.** Linear mixed-effects model of liking ratings by stimulation condition, time, and their interaction. *b*=unstandardized coefficient; *SE*=standard error; CI=confidence interval; Inhib.=inhibitory (cTBS); Excit.=excitatory (iTBS). Excitatory stimulation, T0, and curriculum v1 served as reference categories, so the Condition coefficient is the difference at T0, and the Time coefficients are simple effects within the excitatory condition. The planned contrast is the condition effect at T2 minus the condition effect at T1. The model included random intercepts for participant, melody, and the participant × melody pair. Effect size is Cohen’s *d* (*b* standardized by the model’s residual SD), reported with Cohen’s U3, the proportion of the excitatory condition’s distribution falling below the inhibitory condition’s mean (.50=complete overlap). *pHolm* is reported for the three confirmatory tests (Condition × Time at T1 and T2, and the planned contrast), corrected together as one family; dashes indicate terms that were not part of that family.

| Predictor | *b* | *SE* | *z* | *p* | 95% CI | Effect size | *p* (Holm) |
| --- | --- | --- | --- | --- | --- | --- | --- |
| Intercept | 4.46 | 0.12 | 38.17 | < .001 | [4.23, 4.69] | — | — |
| Condition<br>(Inhibitory vs. Excitatory at T0) | 0.12 | 0.06 | 1.81 | .071 | [-0.01, 0.24] | *d* = 0.14, *U*3 = .56 | — |
| Time<br>(T1 vs. T0, Excitatory) | 0.08 | 0.05 | 1.44 | .149 | [-0.03, 0.18] | *d* = 0.10, *U*3 = .54 | — |
| Time<br>(T2 vs. T0, Excitatory) | 0.23 | 0.05 | 4.22 | < .001 | [0.12, 0.33] | *d* = 0.28, *U*3 = .61 | — |
| Curriculum<br>(v2 vs. v1) | -0.02 | 0.12 | -0.16 | .876 | [-0.25, 0.22] | *d* = -0.02, *U*3 = .49 | — |
| Condition × Time<br>(Inhib. vs. Excit., T1 vs. T0) | -0.07 | 0.08 | -0.94 | .346 | [-0.22, 0.08] | *d* = -0.09, *U*3 = .46 | .346 |
| Condition × Time<br>(Inhib. vs. Excit., T2 vs. T0) | -0.31 | 0.08 | -4.01 | < .001 | [-0.46, -0.16] | *d* = -0.37, *U*3 = .35 | < .001 |
| Planned Contrast |  |  |  |  |  |  |  |
| Difference in Condition Effects<br>(T2 vs. T1) | -0.23 | 0.08 | -3.06 | .002 | [-0.38, -0.08] | *d* = -0.29, *U*3 = .39 | .004 |

### Learning

None of the six condition-performance metrics (three phases × two modalities) reached significance following the application of the Holm correction. For Pitch, the GLMM showed a non-significant trend toward higher acquisition accuracy in logit units under Inhibitory stimulation (b=.12, *p*=.061, pHolm=.184); however, this trend reverses in direction, though remaining non-significant, at both immediate retention (b=-.05, *p*=.482) and delayed retention (b=-.05, *p*=.495) (*see* Table 5). Both conditions demonstrated trial-wise improvement in accuracy, with no condition × trial interaction. This indicates that although participants improved in pitch accuracy as trials were sequentially completed, this improvement was similar across both Excitatory and Inhibitory conditions (*see* Figure 4). A similar directional yet non-significant pattern emerged for rhythm performance. Because rhythm was scored as error (lower values indicating better performance), a positive coefficient reflects worse relative performance under Inhibitory stimulation. Once this is accounted for, rhythm showed the same qualitative pattern as pitch: a small advantage for the inhibitory condition at acquisition (*b*=-.03, *p*=.61, *pHolm*=1.00), that shifted to a small disadvantage at both immediate retention (*b*=.02, *p*=.771, *pHolm*=1.00), and delayed retention (*b*=.07, *p*=.194, *pHolm*=.582). Despite this qualitative pattern, the condition’s main effect was non-significant at acquisition, immediate retention, and delayed retention for Rhythm performance. (*see* Table 5). Similar to Pitch accuracy, percent error in logarithmic units for the acquisition phase was found to decrease significantly across trials, with a significant decelerating quadratic component structure (*b* = .04, *SE* = .01, *z* = 5.85, *p* < .001). Neither trial term interacted significantly with condition (*p*s = .691, .841).

**Table 5.**
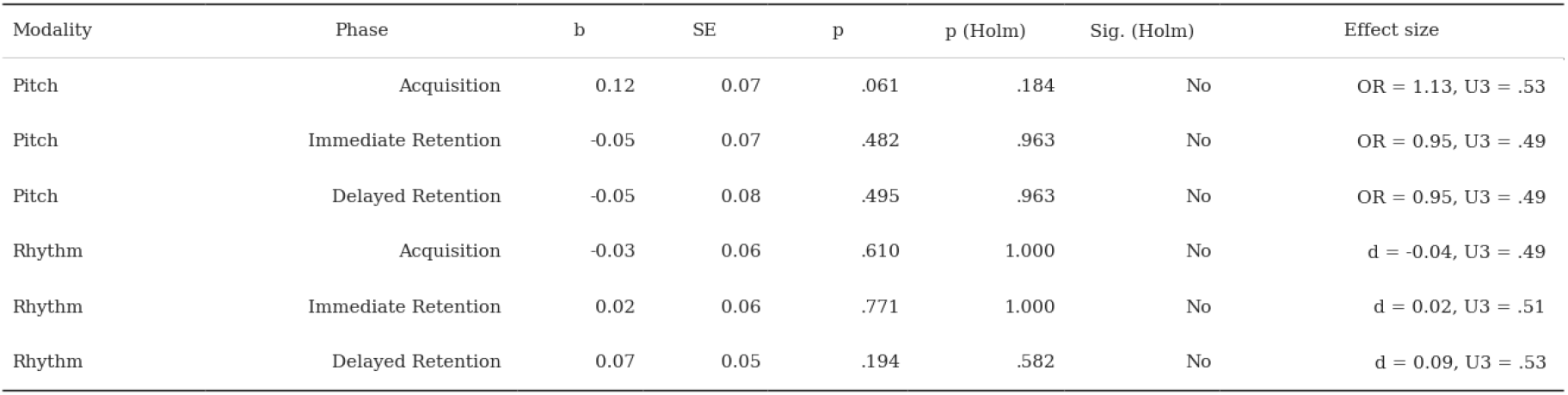
Condition effects on pitch and rhythm performance by learning phase, from the best-fitting model for each phase. Excitatory stimulation (iTBS) was the reference condition, so *b* is inhibitory (cTBS) minus excitatory. Pitch is accuracy (higher=better; logit coefficients from the binomial GLMM) and rhythm is log-transformed relative inter-onset interval error (lower=better; linear mixed-effects models), so a positive coefficient indicates an inhibitory advantage for pitch but a disadvantage for rhythm. Delayed-retention models adjust for immediate-retention performance. Effect size is the odds ratio (OR) for pitch and Cohen’s *d* for rhythm, each with U3, the proportion of the excitatory distribution falling below the inhibitory mean (.50=complete overlap); pitch U3 is computed from a logit-to-*d* conversion (*d* ∼ *b* × 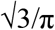; Chinn, 2000). *pHolm* is corrected within each modality (three tests each). Sig.= significant after Holm correction.

| Modality | Phase | b | SE | p | p (Holm) | Sig. (Holm) | Effect size |
| --- | --- | --- | --- | --- | --- | --- | --- |
| Pitch | Acquisition | 0.12 | 0.07 | .061 | .184 | No | OR = 1.13, U3 = .53 |
| Pitch | Immediate Retention | -0.05 | 0.07 | .482 | .963 | No | OR = 0.95, U3 = .49 |
| Pitch | Delayed Retention | -0.05 | 0.08 | .495 | .963 | No | OR = 0.95, U3 = .49 |
| Rhythm | Acquisition | -0.03 | 0.06 | .610 | 1.000 | No | d = -0.04, U3 = .49 |
| Rhythm | Immediate Retention | 0.02 | 0.06 | .771 | 1.000 | No | d = 0.02, U3 = .51 |
| Rhythm | Delayed Retention | 0.07 | 0.05 | .194 | .582 | No | d = 0.09, U3 = .53 |

**Figure 4.**
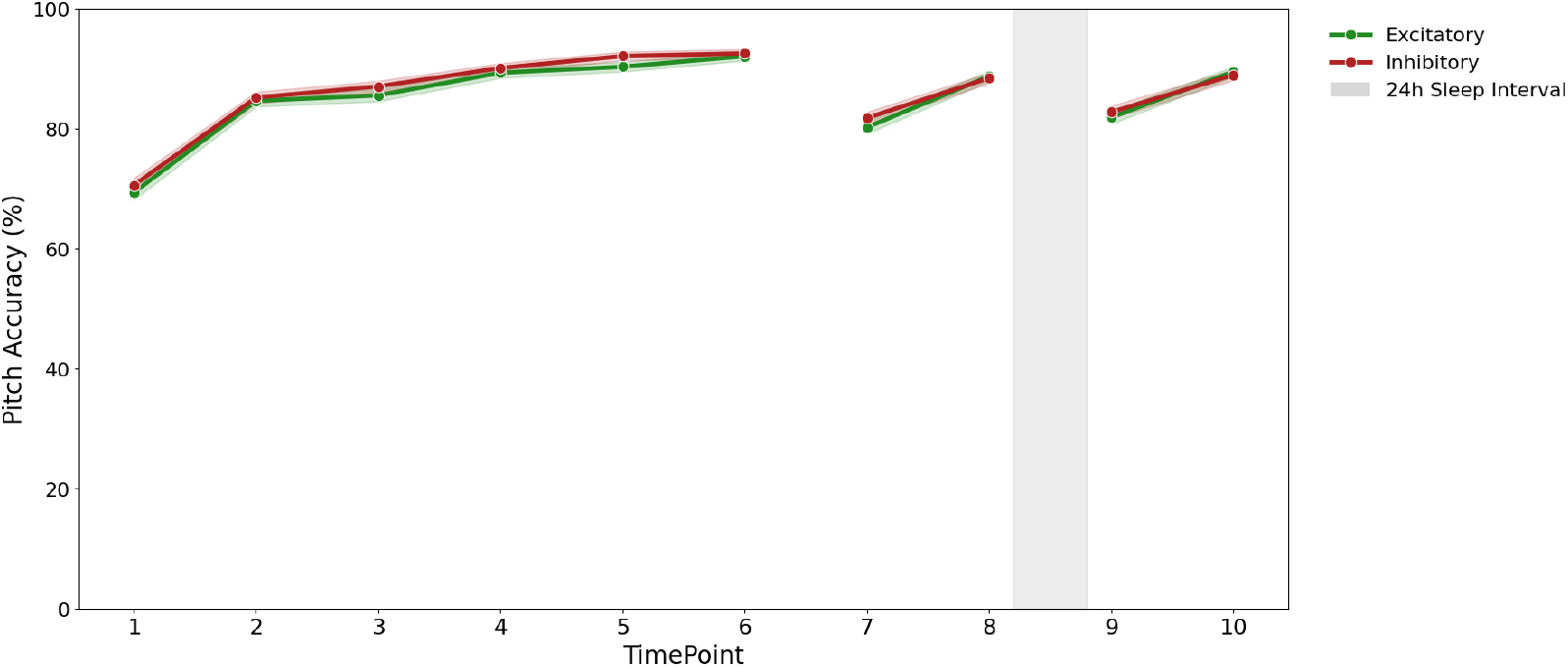
Pitch accuracy across acquisition, immediate retention, and delayed retention by stimulation condition. Lines show mean pitch accuracy (percentage of target notes reproduced correctly) on each trial under excitatory (iTBS) and inhibitory (cTBS) stimulation. Trials 1-6 are acquisition, trials 7-8 immediate retention, and trials 9-10 delayed retention, which took place 24 hours later (gray bar). All trials are shown, including the visually cued trials 2 and 4 that were excluded from the primary acquisition analysis. Shaded bands show ±1 *SE. N*=23.

**Figure 5.**
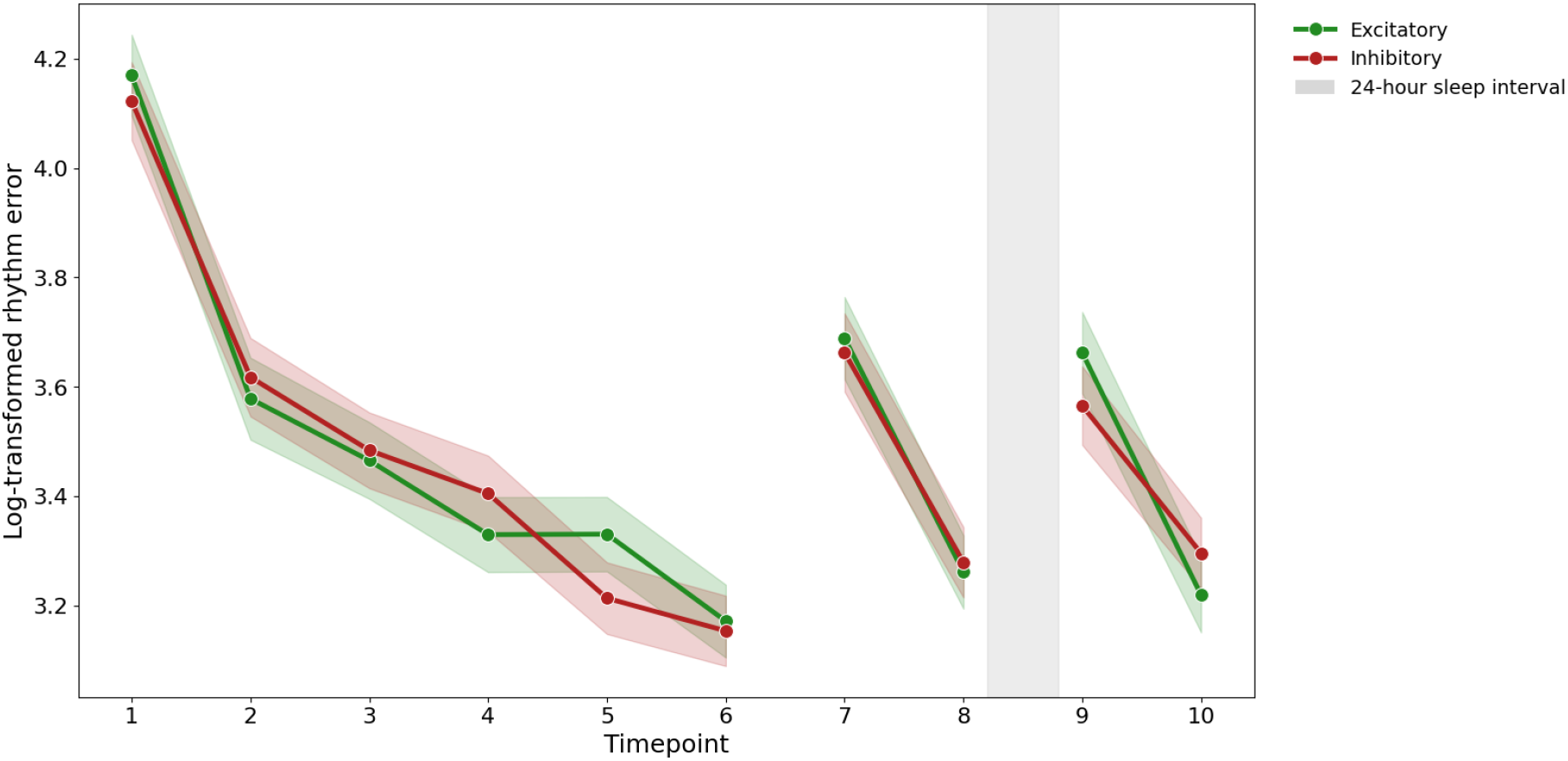
Log-transformed rhythm error across acquisition, immediate retention, and delayed retention by stimulation condition. Lines show mean rhythm error, defined as relative inter-onset interval error (%) transformed with log(1 + x), on each trial under excitatory (iTBS) and inhibitory (cTBS) stimulation. Lower values indicate more accurate reproduction of the melody’s rhythm. Trials 1-6 are acquisition, trials 7-8 are immediate retention, and trials 9-10 are delayed retention, which took place 24 hours later (gray bar). All trials are shown, including the visually cued trials 2 and 4 that were excluded from the primary acquisition analysis. Shaded bands show ±1 *SE. N*=23.

Effect sizes and Cohen’s U3 were therefore examined to characterize the degree of distributional overlap directly. Odds ratios for pitch remained close to 1 (1.13, 0.95, 0.95) and effect sizes for rhythm remained close to 0 (*d*=-.04, .02, .09) across all three phases, and Cohen’s U3, the proportion of the lower-scoring condition’s distribution falling below the higher-scoring condition’s mean, with .50 corresponding to identical distributions, stayed close to the threshold throughout (*see* Table 5). At this degree of overlap, neither condition would be meaningfully over- or under-represented at either extreme of the other’s distribution, as these values describe near-complete overlap between the Excitatory and Inhibitory distributions at every phase of performance in both modalities, consistent with the absence of significant condition effects reported above. However, they do not themselves constitute a formal test of equivalence.

## III. Discussion

Overall, we found that excitatory vs inhibitory theta-burst protocols over the left DLPFC generated a difference in how much participants liked unfamiliar, short piano melodies that they had learned to play. This effect emerged 24 hours following stimulation rather than immediately. We further found that this difference did not extend to how well the melodies were performed during acquisition, immediate retention, or 24 hours after stimulation. Specifically, our principal finding was that liking increased significantly from baseline (T0) to 24 hours following stimulation (T2) in the excitatory iTBS condition but did not change significantly in the inhibitory cTBS condition, and this change differed significantly between conditions, whereas the conditions did not differ significantly between baseline (T0) and ratings collected immediately after TMS (T1). A planned contrast between T1 and T2 confirmed that the divergence arose across the 24-hour interval, which included both the learning task and a period of sleep. Hypothesis (i), that iTBS would increase subjective liking ratings for short piano melodies and cTBS would decrease them, was partly supported: liking increased under iTBS but did not decrease under cTBS, and the difference appeared only at 24 hours.

In contrast, neither pitch nor rhythm accuracy, the behavioral proxy for music playing, differed by condition in any learning phase. Participants, however, improved across acquisition trials in both conditions, indicating that the measures were sensitive to learning. Hypothesis (ii), that learning would follow the same pattern as liking, with melodies whose liking was enhanced under iTBS performed better and melodies under cTBS were performed worse, was not supported.

Therefore, although previous stimulation work has shown that DLPFC theta-burst protocols shift musical pleasure within the session (Mas-Herrero et al., 2018), here we provided evidence that a difference between excitatory and inhibitory protocols can, under certain conditions, be expressed only after a delay. To our knowledge, previous work in the field of reward, music, and learning has not paired a stimulation manipulation of pleasure with a measure of playing the music, thus, including both allowed the present study to ask whether the perception and action steps of the reward cycle are affected together. In our sample, only the perception of hedonic pleasure was affected.

The two studies this study built on each provided guidance to better address whether targeting reward circuitry would have lasting effects on subjective hedonic responses and subsequent early skill acquisition. Mas-Herrero et al. (2018) showed that DLPFC TMS changes musical pleasure immediately: relative to sham, iTBS increased subjective pleasure, physiological arousal, and motivation to purchase music, whereas cTBS decreased these responses. However, that study measured pleasure in the session in which stimulation was delivered, and used long, naturalistic 45s excerpts that were either self- or experimenter-selected to be highly pleasurable. In a follow-up study combining the same protocol with fMRI, Mas-Herrero et al. (2021) showed that stimulation altered striatal responsiveness to hedonic value, and that changes in the ventral striatum, predicted changes in reported pleasure. This follow-up study provided a confirmatory and objective account of the circuitry through which stimulation acts.

Ferreri et al. (2021) had participants rate the pleasure elicited by unfamiliar 20s excerpts during listening and tested their ability to recognize them 24-hours later. Under the placebo condition, excerpts rated as more pleasurable were better remembered, indicating that hedonic responses leave a trace measurable 24 hours later without pharmacological enhancement. Our design differed from both bodies of work in ways that bear on the comparison of our findings. In our study, melodies were 2 to 5 s long and of comparable difficulty so that 20 could be learned in one session. Participants in our design also had two to ten years of musical experience to ensure they would have enough musical background to learn the melodies. Furthermore, we compared iTBS with cTBS directly, rather than with sham, because their opposing effects relative to sham had already been established (Mas-Herrero et al., 2018).

The design differences described above extend earlier TMS work and may explain why the liking effect emerged after a delay rather than immediately. One interpretation concerns the nature of the stimuli. In the Mas-Herrero studies, stimulation was delivered prior to the listening task, thus, its effect on liking presumably depended on how strongly the music engaged the fronto-striatal reward circuitry once it was heard. Musical pleasure derives in large part from the anticipation and resolution of perceptual expectations, where reward prediction errors have been shown to engage the nucleus accumbens (Gold et al., 2019), and the caudate was found to be most engaged during the anticipation of peak pleasure (Salimpoor et al., 2011). The 45s excerpts used by Mas-Herrero et al. (2018, 2021), as well as the 20s excerpts used by Ferreri et al. (2019, 2021) leave time for that temporally unfolding anticipation and resolution to build; two-to-five-second melodies, however, leave little room for it. The melodies were also selected from the highest-rated pilot candidates to replicate neutral to positive pleasure response, which may have restricted the range of liking available to shift. If the circuitry was only weakly engaged when stimulation was delivered, there may have been little activity for the modulation of the hedonic response at T1. A future study which combines melodies graded on hedonic response ability, TMS with a sham condition, and an objective account of activation patterns throughout listening, by means of neuroimaging, would allow these open questions to be addressed.

This interpretation does not contradict the delayed liking effect because T2 did not differ from T1 only by time, but also by a period of learning and of sleep. Between the two assessments, participants completed the learning session, with repeated listening and trial-by-trial feedback, which may have engaged the circuitry more strongly than the T1 ratings did. Sleep-dependent memory consolidation offers a second route by which a small difference at encoding could grow (Brodt et al., 2023). Reward-related processes engaged during encoding have been shown to shape outcomes 24 hours later, an interval that includes a night of sleep (Ferreri et al., 2021). It is therefore plausible that a slight difference in the responsiveness of the reward circuitry was amplified overnight. As exposure, feedback, and sleep were common to both conditions, they cannot explain a condition difference on their own. The stimulation paradigm might have made a difference that was not detectable immediately, but rather provided a slightly higher responsiveness of the reward circuitry under iTBS relative to cTBS, which was then reinforced by learning and consolidation until it was large enough to appear through ratings at T2. This reinforcement concerns the hedonic value assigned to the melodies rather than an ability to play them, and thus need not contradict the null motor performance finding. Thus, this interpretation covers the perception step of the reward cycle, while the action step, production, falls outside what it predicts. Although compatible with the results, this interpretation cannot be tested with the present design. Neural state was not measured, and the contribution of the learning session cannot be separated from that of sleep, as both preceded the T2 rating for every participant. Testing it would require examining the two separately by comparing sessions with and without learning as well as by objectively quantifying sleep with electroencephalography (EEG) proxies of sleep-dependent memory consolidation, such as spindle density and the number of slow oscillations.

This study operationalizes learning as a proxy for the action component of reward, which reinforces the behaviours that follow a hedonic response (Berridge & Kringelbach, 2008), and was designed so that this learning would take place largely within the window of altered excitability.

Whether the learning measured would track the hedonic response was nonetheless an open question, as the evidence that supported it came from a different account of memory: episodic memory, for excerpts that were only heard and then recollected (Ferreri et al., 2021), rather than procedural memory for music production and its retention.

The null finding between conditions and the learning metrics is thus about the action part of the reward cycle as we operationalized it and should not be interpreted as a definite conclusion that music and reward are unrelated to procedural learning and retention. One interpretation is that the circuitry underlying procedural memory and its preferential tagging does not overlap with the circuitry that underlies reward-related episodic memory and its preferential tagging. Work on the DLPFC in procedural learning has linked it to the primary motor cortex with learning-related DLPFC-M1 changes for structured but not random sequences (Cao et al., 2022), and found that stimulation after learning and not before had altered later performance (Galea et al., 2010; Nguyen et al., 2025). Our stimulation paradigm, in accordance with past research on reward and music, preceded learning, and our melodies were simple and short in duration. Therefore, our task may not have been sensitive enough to capture condition effects on learning due to the timing of stimulation and characteristics of the stimuli. This is further supported by the ceiling effects in pitch accuracy, as the melodies may have been simple enough to be learned within a session, but perhaps too simple to elicit preferential tagging. A future study which tests the effects of TMS at different timepoints and with more complex structure is thus warranted to further disentangle the relationship between reward, procedural learning and the manipulation of brain circuitry.

Another interpretation concerns where in the reward cycle reward acts. Playing a melody has its own internal reinforcement, including the visual feedback every participant received and the satisfaction of improving (Badami et al., 2011), which may have driven acquisition regardless of condition (Saemi et al., 2012). This interpretation raises the question of what counts as learning within the reward cycle. In reward literature, the action that a hedonic response reinforces may not be learning to play the stimulus; it may be the reinforcement of its hedonic value, which may be what TMS affects when reward value is minimal. If so, the delayed liking difference may be the learning effect, and our task may have measured a learning process that is dissociable from the manipulation of the reward circuitry. What counts as learning within the reward cycle, whether our manipulation reflects a dissociation between the circuitry supporting liking and that supporting learning, and how each should be measured remain to be determined.

In summary, DLPFC theta-burst stimulation was followed by a difference in musical liking that emerged 24 hours later, which, to our knowledge, has not yet been shown in the field of music perception and reward. Our design informs future studies in two ways. First, melodies short and simple enough to be learned in a single session were still followed by a condition difference in liking, although their limited predictive information should be considered when interpreting the absence of a learning effect. Second, the difference was obtained with BeamF3 scalp-based targeting rather than individual MRI (Mir-Moghtadaei et al., 2015), which makes the paradigm more accessible. Future studies that add a sham condition, graded stimuli, imaging at the time of stimulation, and learning both with and without a delay will help disentangle how reward circuitry contributes to early skill acquisition and consolidation.

## Supporting information

Supplementary Materials (AIC; Table 1-3)

