## Supplementary Materials (AIC; Table 1-3) for "Left-DLPFC theta-burst stimulation produces delayed effects on musical pleasure without affecting piano-melody learning"

| Phase | Raw AIC | Logit (corrected) AIC | Δ AIC | Better-Fitting Model |
|---|---|---|---|---|
| Acquisition | 30857.75 | 19894.08 | 10963.67 | Logit |
| Immediate retention | 7565.00 | 4603.28 | 2961.72 | Logit |
| Delayed retention | 7382.49 | 4177.90 | 3204.59 | Logit |

**Table 1. Fit comparison of linear mixed-effects models for pitch performance using raw and logit-transformed percentages, by phase.** AIC=Akaike information criterion; lower values indicate better fit. Logit AIC values are Jacobian-corrected so that they are comparable with the AIC of models fit to raw percentages. ΔAIC=Raw AIC-Logit AIC.

| Phase | GLM AIC | GLMM AIC | Δ AIC | Better-Fitting Model |
|---|---|---|---|---|
| Acquisition | 14197.88 | 9014.78 | 5183.10 | GLMM |
| Immediate retention | 3386.44 | 2110.97 | 1275.47 | GLMM |
| Delayed retention | 2310.75 | 1967.80 | 342.95 | GLMM |

**Table 2. Fit comparison of generalized linear and generalized linear mixed-effects models for pitch performance, by phase.** GLM= generalized linear model without random effects; GLMM=generalized linear mixed-effects model with random effects for participant and melody. AIC=Akaike information criterion; lower values indicate better fit. ΔAIC=GLM AIC- GLMM AIC.

| Phase | Linear AIC | Quadratic AIC | Δ AIC | Better-Fitting Model |
|---|---|---|---|---|
| Acquisition | 9956.36 | 9895.75 | 60.61 | Quadratic |

**Table 3. Fit comparison of linear and quadratic trial curves for log-transformed rhythm error during acquisition.** Rhythm error is the relative inter-onset interval error (%), transformed with log(1+x). Linear and quadratic refer to the form of the centered trial term, each entered with its interaction with stimulation condition. AIC=Akaike information criterion; lower values indicate better fit. ΔAIC=Linear AIC- Quadratic AIC
